# Making sense of virus size and the tradeoffs shaping viral fitness

**DOI:** 10.1101/2020.04.01.020628

**Authors:** Kyle F. Edwards, Grieg F. Steward, Christopher R. Schvarcz

## Abstract

Viruses span an impressive size range, with genome length varying more than a thousandfold and capsid volume nearly a millionfold. Physical constraints suggest that smaller viruses may have multiple fitness advantages, because a greater number of viral offspring can be created from limited host resources, and because smaller particles diffuse to encounter new hosts more rapidly. At the same time, a larger genome size allows for numerous additional functions that may increase fitness, such as better control of replication, transcription, translation, and host metabolism, and neutralization of host defenses. However, it is unclear whether important viral traits correlate with size, and whether this causes size to vary among host types or environmental contexts. Here we focus on viruses of aquatic unicellular organisms, which exhibit the greatest known range of virus size. We develop and synthesize theory, and analyze data where available, to consider how size affects the primary components of viral fitness. We suggest that the costs of larger size (lower burst size and diffusivity) are mitigated by the role of a larger genome in increasing infection efficiency, broadening host range, and potentially increasing attachment success and decreasing decay rate. These countervailing selective pressures may explain why such a breadth of sizes exist and can even coexist when infecting the same host populations. We argue that oligotrophic environments may be particularly enriched in unusually large or “giant” viruses, because environments with diverse, resource-limited phagotrophic eukaryotes at persistently low concentrations may select for broader host range, better control of host metabolism, lower decay rate, and a physical size that mimics bacterial prey. Finally, we describe areas where further research is needed to understand the ecology and evolution of viral size diversity.

## Introduction

Viruses are ubiquitous and abundant parasites that influence individual health, population and community dynamics, evolution, and biogeochemistry, across the tree of life. By nature, viruses are smaller than the cells they infect, but the range of virus sizes is nonetheless substantial, with lengths of viral particles (virions) varying from 17 nm to ~1.5 μm, and genome size varying from ~1 kb to 2.5 Mb (Campillo-Balderas et al. 2015). The largest ‘giant’ viruses have primarily been isolated from unicellular protists (Campillo-Balderas et al. 2015, Wilhelm et al. 2017), although there is metagenomic evidence for ‘megaphages’ of prokaryotes with genomes up to 716 kb (Devoto et al. 2019, Al-Shayeb et al. 2020), and a chaetognath appears to be infected by viruses 1.25 μm in length (Shinn and Bullard 2018). In contrast, the known viruses of plants and fungi have genomes < 30 kb (Campillo-Balderas et al. 2015). At a finer phylogenetic scale, particular species or strains of prokaryotes and eukaryotes can be infected by viruses of very different size. For example, the marine bacterium *Cellulophaga baltica* is infected by phages ranging from 6.5 to 242 kb (Holmfeldt et al. 2007) and the marine dinoflagellate *Heterocapsa circularisquama* is infected by a 4.4 kb ssRNA virus and a 365 kb dsDNA virus (Tomaru et al. 2009).

For cellular life, body size is a ‘master trait’ that influences numerous organismal properties, such as metabolic rate, nutrient uptake affinity, predator-prey linkages, and population growth rate (Finkel 2001, Brown et al. 2004, Fuchs and Franks 2010, Edwards et al. 2012). The substantial variation in virus size raises the question of whether size plays a similar central role in virus ecology and evolution (Record et al. 2016). For example, are there general relationships between virus size and key viral traits? Do size-related tradeoffs lead to selection for different sizes of viruses infecting different kinds of hosts, or under different environmental conditions? How do viruses of different size coexist when infecting the same host population? There are straightforward physical reasons why being smaller should be advantageous for a virus: smaller particles should encounter hosts faster due to greater diffusivity, at least in aquatic systems or aqueous microenvironments, and limited host resources during infection can be partitioned among a greater number of ‘offspring’. The existence of a spectrum of virus sizes implies that the costs of increased size can be offset by countervailing benefits. Virion size and genome size are tightly correlated (Cui et al. 2014), and benefits of increased size likely derive from the functions encoded by viral genes, including better control of attachment to the host, replication, transcription, translation, and host metabolism; strategies countering host antiviral defenses; and repair of damaged viral nucleic acids (Sharon et al. 2011, Samson et al. 2013, Fischer et al. 2014, Koonin and Yutin 2019, Mendoza et al. 2019). These functions could increase the probability of successful infection, the number of virions produced per infection, the range of hosts than can be successfully infected, and/or the persistence of virions in the extracellular environment.

In this study, we focus on the question of how virus size affects key viral traits, and how these traits affect viral fitness. We develop and summarize relevant theory, synthesize and analyze available data, and outline major knowledge gaps and future research directions. We focus primarily on viruses that infect aquatic microbes (unicellular prokaryotes and eukaryotes), as these viruses are known to vary greatly in size and have been relatively well-studied in culture. However, most of the concepts we develop should be useful for understanding viruses in general.

### Theory for how viral fitness is determined by key viral traits

In order to connect a metric of fitness to virus traits (burst size, latent period, contact rate, decay rate, etc.) we imagine a lytic virus population that may compete for hosts with one or more additional virus populations. Using a simple model of virus-microbe population dynamics, at steady state the density of the limiting resource, uninfected cells, is 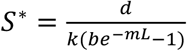 (eqn. 1; Appendix 1). Here *S* is the density of uninfected cells, *m* is the host mortality rate from causes other than viral infection, *b* is the burst size (new virions produced per infection), *L* is the latent period, *d* is the viral decay rate, and *k* is the effective adsorption rate—the rate at which successful new infections are formed. The parameter *k* can be decomposed into subprocesses, and here we define *k* = *caw*, where *c* is the contact rate at which host and virus encounter each other, *a* is the attachment efficiency (probability that encounter leads to successful attachment), and *w* is the probability that attachment leads to a successful infection (eventual lysis of the host, releasing new virions). The distinction between contact rate and attachment efficiency will be important when considering the role of virus size.

The quantity *S*\* is the uninfected host density at which growth of the viral population balances the decay rate. Therefore, this is also the host density threshold required for persistence of a viral population, and if the host density is initially greater it will be cropped down to this level at steady state. *S*\* can be used as a measure of viral fitness because if there are two viral strains, and 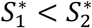, then strain 1 will drive the host to a lower density and competitively exclude the other strain at equilibrium (Tilman 1982). It is more intuitive to consider the inverse of *S*\* as a metric of fitness, because a smaller S* equates to greater competitive ability; the inverse of *S*\* is 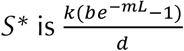. This quantity is the average net number of new virions produced per infection (*be*^−mL^ − 1), scaled by the rate of successfully encountering a new host (*k*) relative to the rate of ‘dying’ while waiting to encounter a new host (*d*). This analysis assumes that host-virus dynamics reach a steady-state attractor, which may not be true, but the *S*\* quantity is still a useful index of competitive ability. Furthermore, in a simpler model with no latent period it can be shown that *S*\* predicts the winner in competition even if populations fluctuate (Appendix 2). This analysis relates traits to fitness for lytic viruses, which we focus on in this study due to a greater accumulation of relevant trait data, but potential effects of size on viruses with temperate strategies will be discussed as well.

In Appendix 3 we extend this analysis to ask under what conditions a broader host range is selected for. The main result is that fitness (measured in terms of competitive outcomes) is proportional to host range (measured as the number of host strains that can be infected). This means that a generalist virus and a specialist virus will have similar fitness when the cost of generalism is directly proportional to host range breadth. For example, if a generalist virus can infect twice as many strains as specialist viruses, but has an adsorption rate that is 50% lower on each strain, it will be competitively equivalent to the specialists. If the cost of generalism is lower then generalism will be favored, and vice versa. These results are derived from a simple model but they allow us to quantify, as a first approximation, how tradeoffs involving host range and other viral traits may affect selection on virus size.

### How does virus size affect contact rate and attachment to host cells?

We now consider key viral traits individually, to outline expectations for how virus size may affect each trait and analyze relevant data where it is available. Adsorption rate is important for viral fitness (eqn. 1) because it determines the rate at which new infections can be established, as well as the time a viral particle spends in the extracellular environment where it may be exposed to UV radiation, adsorption to non-host material, ingestion, etc. (Suttle and Chen 1992, Noble and Fuhrman 1997). As described above, it is useful to separate the adsorption rate into the contact rate *c* (per capita rate at which hosts and viruses encounter each other) and the attachment efficiency *a* (probability of successful attachment to the host). In theory the contact rate will depend on physical processes of Brownian motion, advection, and turbulence, while attachment efficiency will be a function of host receptor availability, affinity of the receptor for the virus, and mechanisms such as reversible binding by viral fibers that keep the virus from diffusing away from the host (Schwartz 1976, Wickham et al. 1990, Storms and Sauvageau 2015).

Physical theory for contact rate typically starts by asking what the rate would be if viruses relied solely on molecular diffusion to encounter their hosts, and if all viruses that contact the host are adsorbed. Under pure diffusion the contact rate is predicted to be:

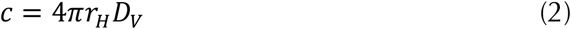

where *r*_H_ is host radius and *D*_V_ is the diffusivity of the virus (Murray and Jackson 1992). The diffusivity of a spherical virus is predicted to be:

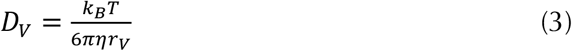

where *k*_B_ is the Boltzmann constant, *T* is temperature, *η* is dynamic viscosity of water, and *r_V_* is the virion radius. Therefore, the diffusion-limited contact rate is predicted to be inversely proportional to virus diameter, which is a substantial fitness cost of increasing size.

The diffusivity of viruses due to Brownian motion is low enough that contact rates could be increased considerably by processes that create fluid motion relative to the host cell. Rates of diffusion can be enhanced by advective flow arising from host motility, feeding currents, or host sinking, and diffusion can also be enhanced by turbulence, which causes shear around the host cell (Murray and Jackson 1992). Host motility, feeding currents, sinking, and turbulence can also lead to the host cell encountering the virus by direct interception (i.e., without the aid of Brownian motion), which is the mechanism by which small flagellates are thought to encounter immotile prey (Shimeta 1993, Kiørboe 2008).

The results in Appendix 4 show that diffusion enhanced by advection is the primary mechanism that could significantly increase virus contact rates beyond the ‘pure diffusion’ scenario (eqn. 2), and so we consider here how that mechanism depends on virus size. An approximate formula for contact rate when advection enhances diffusion is:

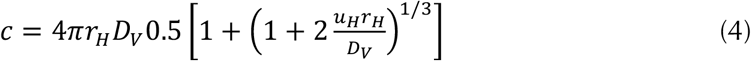

where *u_H_* is the velocity of the host relative to the surrounding water (Murray and Jackson 1992). Because the formula includes the term 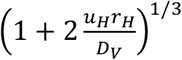, the enhancement of contact rates due to advection is greater as *u_H_* and *r_H_* increase, and also as *r_V_* increases (because *D*_V_ is in the denominator, and *D*_V_ ~ 1/*r*_V_). In other words, host motility matters more for bigger hosts, for hosts that swim faster, and for bigger, less diffusive viruses. These effects are visualized in Fig. 1A, which shows that contact rate always declines with virus size, but the penalty for large size is slightly less when hosts are motile. The effect of host motility on contact rates ranges from modest (~2-fold increase for a 20 nm virus infecting a 1 μm host swimming at 30 μm s^−1^) to large (>10-fold increase for a 300 nm virus infecting a 20 μm host swimming at 250 μm s^−1^). Langlois et al. (2009) use numerical simulation to show that the effect of swimming on diffusion may be underestimated by the formulas used here, but the effect is only ~2-fold for the relevant particle sizes and swimming speeds.

**Figure 1.**
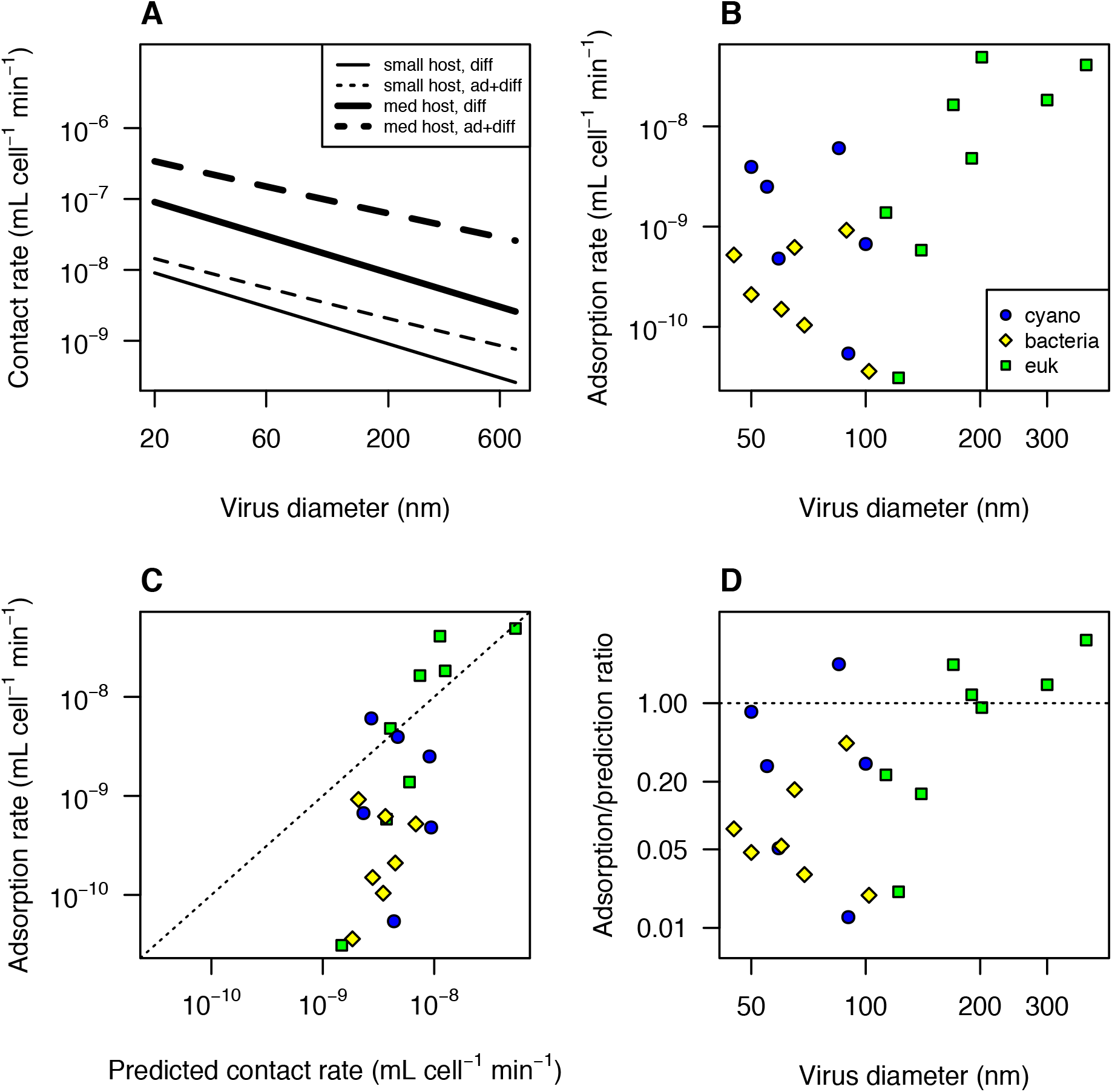
Model predictions and observations of contact rate and adsorption rate. (A) Predicted contact rate as a function of virus size. The ‘small host’ lines correspond to a host with diameter 1 μm and swimming speed 10 μm s^−1^, and the ‘medium host’ lines correspond to a host with diameter 10 μm and swimming speed 100 μm s^−1^. The solid lines are the pure diffusion prediction (no advection), and the dashed lines are for swimming hosts (advection+diffusion). (B) Observed adsorption rates for viruses of aquatic bacteria, phytoplankton, and the protist *Cafeteria roenbergensis* (Table S1). (C) Observed rates vs. theoretical predictions described in Methods. (D) Observed adsorption rate, relative to the theoretical maximum contact rate, as a function of virus size. ‘cyano’ = virus infecting cyanobacterium, ‘bacteria’ = virus infecting heterotrophic bacterium, ‘euk’ = virus infecting unicellular eukaryote.

Although Brownian motion, potentially enhanced by advection, is expected to drive contact rates, it is possible that direct interception is important for particularly large viruses encountering hosts that generate a strong current (Figs. S1−2). The simple model of interception based on Stokes flow may underestimate particle contact rates (Langlois et al. 2009), and if interception is great enough then ‘giant’ viruses could have greater contact rates than slightly smaller viruses (Figs. S1−2). Therefore, a better understanding of the physics of particle encounter will be important for understanding virus ecology and size evolution, in addition to understanding predator-prey dynamics among microbes.

We have compiled published data on the adsorption rates of viruses of aquatic microbes (cyanobacteria, heterotrophic bacteria, eukaryotic phytoplankton, heterotrophic protists; Table S1) to ask whether virus size has any relation to contact rate or attachment efficiency. There is a tendency for adsorption rate to be greater for larger viruses (Fig. 1B), but this may be due to larger viruses having hosts that are larger, more motile, or both. We therefore used eqns. 2−4 to ask how observed adsorption rates compare to the theoretical maximum (Fig. 1C). Predictions and observations are positively correlated (r = 0.68), but many of the observations are 10-100x lower than predicted. This is consistent with a previous analysis of adsorption rates of phages (Talmy et al. 2019) and could be due to sparse host receptors, a low binding affinity of the virus ligand to the host receptor, or a lack of mechanisms for keeping the virus from diffusing away from the host before irreversible binding to the receptor occurs (Storms and Sauvageau 2015). Several of the large eukaryotic viruses have adsorption rates that are higher than the prediction based on Brownian motion alone (eqn. 2), but accounting for host swimming (eqn. 4) brings them closer to the 1:1 line (Fig. S3). However, a number of the phages also have motile hosts, and including reasonable numbers for host swimming speed moves them further below the 1:1 line (Fig. S3).

Fig. 1D shows the proximity of adsorption rate to the theoretical maximum as a function of virus size. Although the sample size is limited, it is noteworthy that the five largest viruses are all fairly close to the theoretical maximum. It is possible that some of the functions encoded in larger genomes increase attachment efficiency, such as the synthesis of proteins that aid attachment to host glycans (Rodrigues et al. 2015), or a greater diversity of proteins for binding host receptors (Schwarzer et al. 2012). If attachment efficiency is promoted strongly by certain genes, this could outweigh the reduction in diffusivity associated with larger size, increasing the actual adsorption rate. If a virus is large enough to induce phagocytosis this could also potentially increase encounter efficiency relative to other mechanisms of entry, and some of the largest viruses have been shown to enter their amoeba hosts via phagocytosis (Rodrigues et al. 2016). If phagocytosis is in fact a more efficient entry mechanism then large size could be selected for, in order to induce phagocytosis (Rodrigues et al. 2016). Protists that eat bacteria-sized prey are known to ingest larger prey at higher rates, which could be due to differences in contact rates or size preferences during ingestion (Chrzanowski and Šimek 1990, Holen and Boraas 1991, Šimek and Chrzanowski 1992, Epstein and Shiaris 1992). Additional measurements of adsorption rates for viruses across the full size spectrum will be needed to test whether larger size is on average an advantage or disadvantage, and whether rates of successful encounter and infection differ for viruses that enter by phagocytosis.

### How does virus size affect viral production during infection?

Burst size and latent period of a lytic virus are determined by the rate at which new virions are created during an infection and the timing of cell lysis (You et al. 2002). The production of virions may decline as host resources are depleted, and the timing of cell lysis may evolve in response to intracellular conditions, host density, and other factors (Wang et al. 1996, Abedon et al. 2003). A previous analysis of phytoplankton viruses showed that burst size of dsDNA viruses may be limited by the host resources used in virus genome replication, with lysis occurring once those resources are exhausted (Edwards and Steward 2018). By contrast, small ssDNA and ssRNA viruses that infect large hosts may maximize fitness by lysing the host before those resources are exhausted (Edwards and Steward 2018).

In light of these results, we focus here on the role of viral genome size in constraining burst size and the rate of viral replication. Eqn. 1 shows that burst size and latent period are expected to play a large role in virus fitness, and therefore viral size evolution may be driven in part by its effects on these life history parameters. We previously showed that burst size is correlated with the host:virus genome size ratio (Edwards and Steward 2018), and this relationship can be decomposed into the effects of host genome size and virus genome size. Of the total variation in burst size across phytoplankton viruses, 48% is explained by host and virus genome sizes in combination, with a partial R^2^ of 30% for host genome size and a partial R^2^ of 14% for virus genome size (4% of variation cannot be uniquely attributed to either predictor because host and virus genome sizes are partially correlated). Although burst size tends to decline for larger viruses, the effect of virus genome size is less than proportional, i.e., a tenfold increase in genome size leads to a less than tenfold decrease in burst size (Fig. 2A). The estimated slope for virus genome size is −0.52 (95% CI = [-0.225, −0.911]) when host and virus taxonomy are included as random effects, or a slope of −0.3 (95% CI = [-0.11, −0.55]) when host and virus taxonomy are not included. This means that a tenfold increase in virus genome size would be expected to reduce burst size by a factor of 1/10^−0.52^ = 3.3 or 1/10^−0.3^ = 2.0.

**Figure 2.**
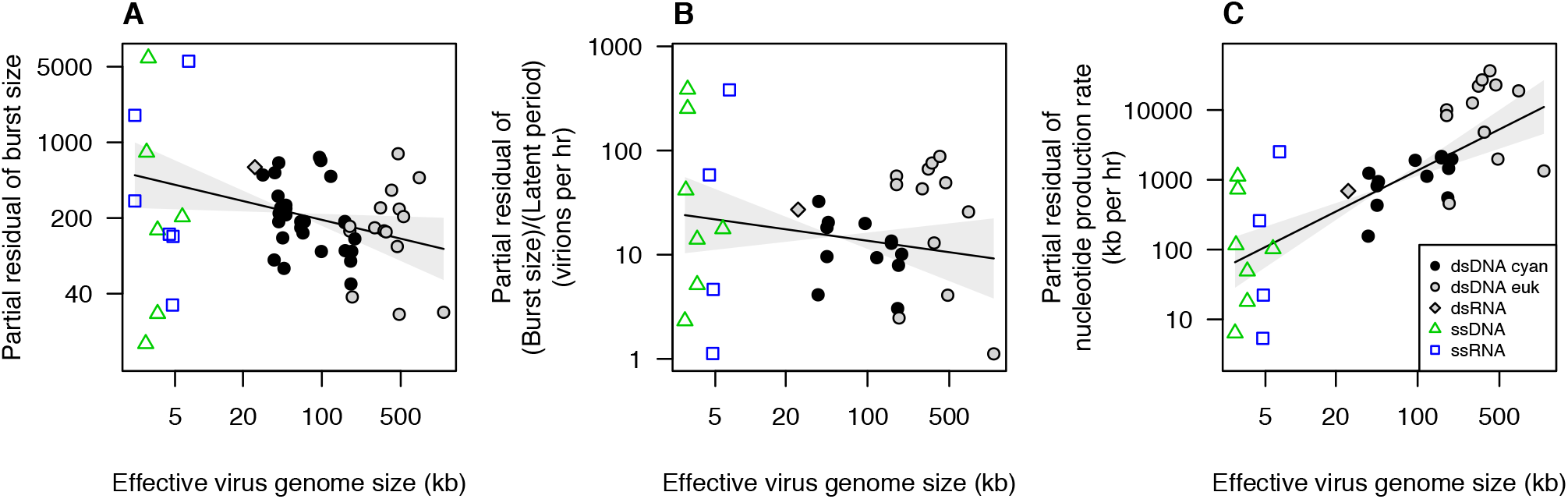
Burst size and production rates as a function of virus genome size for viruses of phytoplankton and the protist *Cafeteria roenbergensis.* (A) Burst size vs. virus genome size. Partial residual burst size is plotted, which removes variation in burst size explained by host genome size. Effective virus genome size is plotted, which divides genome size by 2 for single-stranded viruses. (B) Virion production rate, (burst size)/(latent period), vs. virus genome size. Partial residual production rate is plotted, which removes variation explained by host growth rate. (C) Nucleotide production rate (kb per hr) vs. virus genome size. Partial residual production rate is plotted, which removes variation explained by host growth rate. Plotted lines are fitted smoothers ± 95% CI from generalized additive mixed models.

A less-than-proportional relationship between virus genome size and burst size suggests that larger viruses are producing more total viral material per infection, which could happen if viruses with larger genomes are better at extracting resources from their hosts, better at maintaining metabolic processes that fuel replication, more efficient at transcription or translation, etc. For another perspective on the same processes we can consider how quickly new virions are produced during an infection. All else equal, we would expect that larger virions take longer to construct, due to rate limitation by protein elongation, supply of amino acids or dNTPs, or other processes (You et al. 2002, Birch et al. 2012). The data for phytoplankton viruses exhibit a weak trend of virion production rate declining with virus size (F_1,36_ = 2.3, p = 0.14; Fig 2B), which is consistent with a penalty for larger size that is not directly proportional to size. Finally, rather than looking at production rate on a per virion basis we can consider production rate on a per nucleotide basis, to quantify the total rate at which viral nucleotides are produced during an infection. Nucleotide production rate increases strongly with viral size, with the largest viruses on average producing viral nucleotides ~100x faster than the smallest viruses (Fig. 2C; F_1,11_ = 6.5, p = 0.027). In combination with Fig. 2A-B, this argues that larger size does incur a cost of producing fewer offspring per infection, but that the cost is partially mitigated by a more effective infection process facilitated by the functions encoded in larger genomes. An important question for future research is how resource limitation affects these relationships. Nutrient or light limitation have been shown to reduce burst size and lengthen latent period (Wilson et al. 1996, Maat and Brussard 2016, Thamatrakoln et al. 2019), but the data analyzed here are from experiments under resource-replete conditions. It is possible that the advantage of large genome size is greater under resource limitation, due to a greater control of rate-limiting metabolic reactions.

### Do larger viruses have broader host ranges?

Relatively large viruses could be favored in competition with smaller viruses if a larger genome size is associated with a broader host range (Appendix 3; Chow and Suttle 2015). There is substantial evidence that the ability to attach to host cells plays a major role in defining viral host range (Tétart et al. 1996, Tarutani et al. 2006, Stoddard et al. 2007, Lin et al. 2012, Le et al. 2013), and laboratory experiments often find that hosts evolve resistance by limiting attachment (Lenski 1988, Stoddard et al. 2007). Therefore, virus size may be correlated with host range if having more genes facilitates attachment to a broader range of receptors. For example, the large myovirus phi92 possesses a ‘Swiss army knife’ of multiple tail fibers and/or spikes that appears to facilitate a relatively broad host range encompassing diverse *Escherichia coli* and *Salmonella* strains (Schwarzer et al. 2012). In addition, if the largest viruses are typically ingested by their hosts then they may have a broader host range than smaller non-ingested viruses, due to the fact that ingestion of prey tends to be a less specific interaction than ligand-receptor binding.

Although attachment is known to be important in defining host range, in some cases viruses can attach with equal or lesser efficiency to related strains or taxa that do not yield productive infections (Samimi and Drews 1978, Thomas et al. 2011, Yau et al. 2018). Presumably the infections were not productive in these cases because of many processes downstream of attachment that can limit infection success, such as successful entry into the host, intracellular host defense mechanisms, and effective meshing with host replication and translation machinery (Samson et al. 2013). Furthermore, attaching to a relatively narrow range of cell types in the environment can be adaptive if broader attachment results in a loss of virions to hosts that cannot be infected as productively (Heineman et al. 2008). Therefore, the ability to create productive infections once attached may play a large role in defining host range, and viruses with more genes that provide greater autonomy during infection, via control of replication, transcription, translation, or metabolism, or by evading host defenses, may achieve productive infections in a broader range of hosts.

To assess whether there is an association between genome size and host range we reanalyzed data from two studies of marine phages. Holmfeldt and colleagues characterized a taxonomically diverse collection of 40 phages isolated on 21 strains of the marine bacterium *Cellulophaga baltica* (Holmfeldt et al. 2007, Holmfeldt et al. 2016, Sulcius and Holmfeldt 2016). Across these phages there is an overall positive correlation between host range and genome size, if host range is quantified as the proportion of 21 *Cellulophaga* strains infected (Fig. 3A; r = 0.49, p = 0.001). We also quantified host range in a way that incorporates phylogenetic relatedness of the host strains, using the summed branch lengths of the infected strains, calculated from a phylogeny estimated from ribosomal internal transcribed spacer (ITS) sequences; this yielded similar results (results not shown). A positive relationship also appears to occur within phage families, with larger isolates of myovirus, siphovirus, and podovirus having a greater host range than smaller isolates within the same groups (Fig. 3A). However, when phage family and phage genus (as classified by Holmfeldt et al. [2016]) are both included as a random effects in a mixed model, in order to account for phylogenetic non-independence in host range (Felsenstein 1985), the effect of host genome size is less clear (χ_1_ = 2.45, p = 0.12). This suggests that an even greater phylogenetic diversity of viruses may be needed to robustly test such relationships using a comparative approach. Wichels et al. (1998) characterized 22 phages from the North Sea that infect the bacteria *Pseudoalteromonas*. Across the phages in this study there is also a positive correlation between genome size and host range (r = 0.79, p < 0.001), and evidence for such a relationship remains after taxonomic random effects are included in a generalized additive mixed model (χ_1.2_ = 7.2, p = 0.007; taxonomic terms include family, morphotype, and species as defined by the authors).

In sum, it may be the case that viruses with larger genomes tend to infect a broader range of hosts, and future analyses from diverse host-virus systems would help test the generality of this pattern. At the same time, it is noteworthy that among *Cellulophaga* phages the smallest phage family, the Microviridae, exhibit relatively broad host range on average (Fig. 3A). A large study of *Vibrio* phages also found that small phages, which the authors classified as Autolykiviridae, had broader host ranges than larger Caudovirales (Kauffman et al. 2018). Future work incorporating a quantitative metric of viral fitness on each host strain would help test whether small, broad-range viruses suffer a cost of lower fitness on each individual host strain (Jover et al. 2013, Record et al. 2016). Tradeoffs affecting viral traits, including those related to size and those orthogonal to size, are likely multidimensional (Goldhill and Turner 2014), and therefore it will be important to measure multiple traits on a diversity of viruses to better understand the constraints on viral evolution.

**Figure 3.**
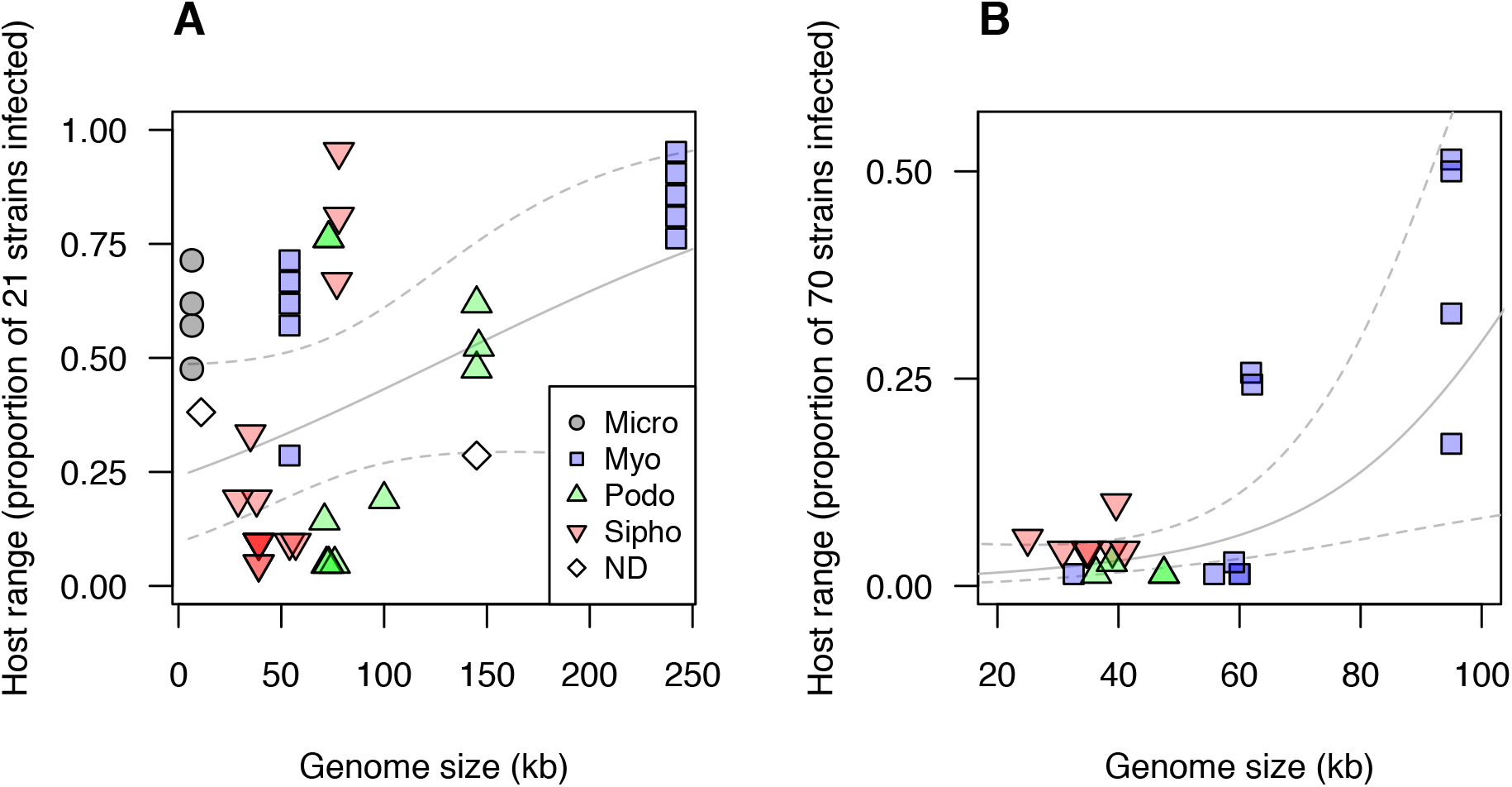
Host range vs. genome size in marine bacteriophages. (A) Host range (proportion of 21 strains infected) vs. genome size for *Cellulophaga baltica* phages (Sulcius and Holmfeldt 2016). (B) Host range (proportion of 70 strains infected) vs. genome size for *Pseudoalteromonas* phages (Wichels et al. 1998). Lines depict fitted smoothers and 95% CI from generalized additive mixed models that include taxonomic random effects to account for related viruses having similar host ranges. ‘Micro’ = Microviridae, ‘Myo’ = Myoviridae, ‘Podo’, = Podoviridae, ‘Sipho’ = Siphoviridae, ‘ND’ = taxonomy not determined.

### Are larger viruses more persistent in the environment?

The rate at which free virions are lost from a viral population is as important for fitness as adsorption rate, burst size, or host range (eqn. 1). However, the effects of virus size on loss rates are poorly known. Decay of phage infectivity in marine systems has been shown to be influenced by sunlight, adsorption to particles, high molecular weight dissolved material such as enzymes, and ingestion by protists (Suttle and Chen 1992, Noble and Furhman 1997). Although a number of studies have estimated decay rates and how they vary across environmental gradients, we are not aware of studies that look at whether these rates vary systematically with size. Heldal and Bratbaak (1991) noted that viruses > 60 nm disappeared more slowly when viral production was halted with cyanide, but they presented no quantitative data.

The physical forces that affect virion stability likely vary with size. For viruses with larger double-stranded genomes the capsid can be highly pressurized due to the dense packaging of negatively charged, dehydrated, curved nucleic acids (Purohit et al. 2003, Li et al. 2008, Molineux and Panja 2013). In a comparative study of coliphages, De Paepe and Taddei (2006) found that phages with a faster multiplication rate in culture had a faster decay rate as well. Faster decay was also associated with a higher nucleotide packaging density and a lower surfacic mass (capsid molecular weight per capsid surface area), suggesting that greater pressure makes capsids less stable, and this can be partially mitigated by increased capsid thickness. A re-analysis of their data shows that virus diameter is also negatively correlated with decay rate (r = −0.64; mixed model with virus family random effect − F_1,13_ = 9.1, p = 0.01; Fig. 4A), which could be due to larger phages having lower packaging density, higher surfacic mass, or other causes.

**Figure 4.**
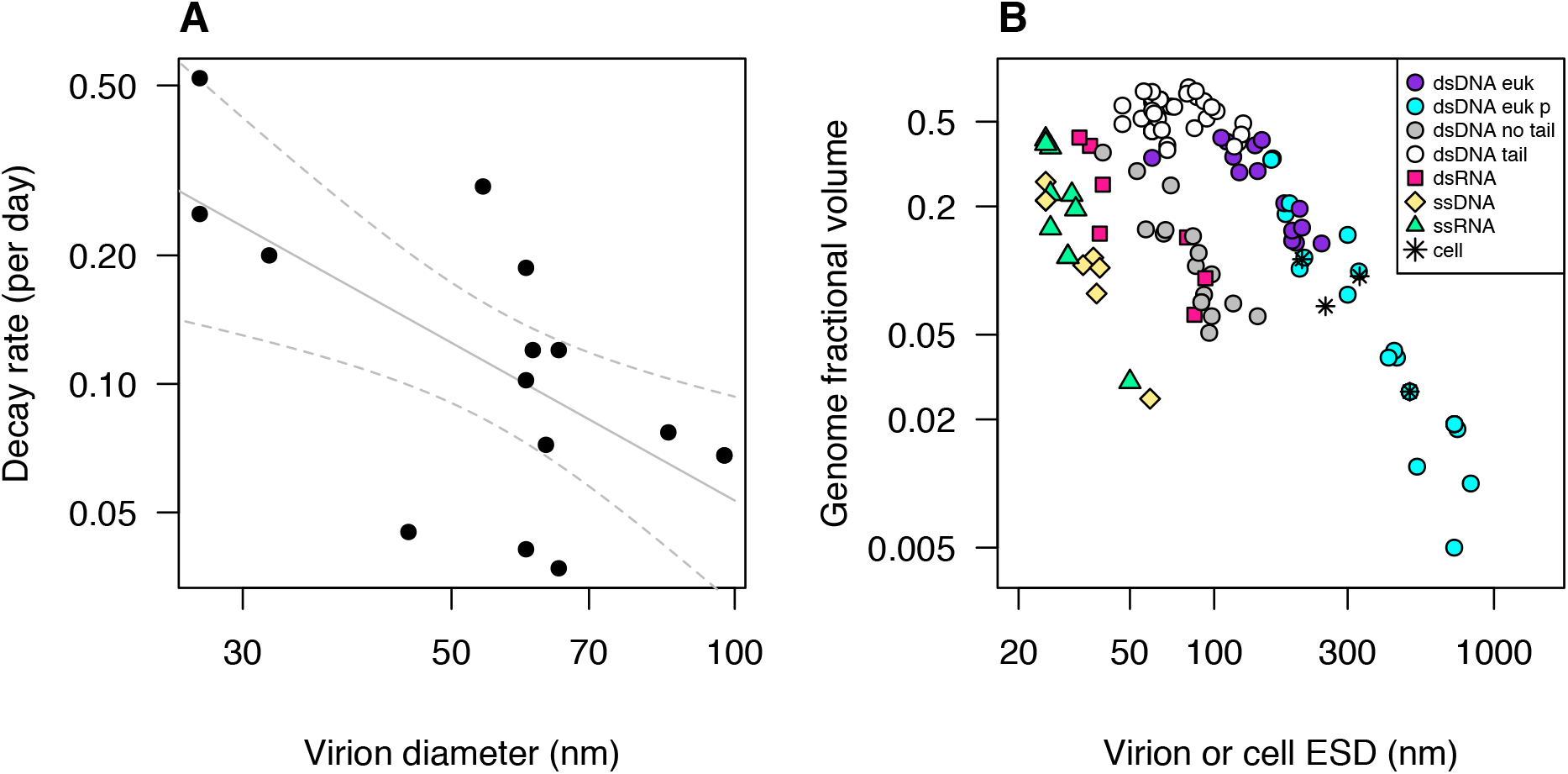
(A) Decay rate vs. virion diameter, for phages that infect *E. coli* (data from Table 1 in De Paepe and Taddei 2006). Lines are linear regression fit ± 95% CI. (B) Genome fractional volume vs. equivalent spherical diameter, for viruses that infect unicellular organisms, and for four representative cellular organisms. Genome fractional volume is the estimated nucleic acid volume divided by total virion volume. Cell or virion volumes were estimated from reported outer dimensions (including outer envelope, if present, but excluding tails or fibrils) using volume formulae for simplified approximate shapes as indicated. Nucleic acid volume is estimated as volume of a cylinder with diameter being 2.37 nm for double-stranded nucleic acid and 1.19 nm for single-stranded and 0.34 nm per nt or bp. ‘dsDNA euk’ = dsDNA viruses infecting non-phagotrophic eukaryotes, ‘dsDNA euk p’ = dsDNA viruses infecting phagotrophic eukaryotes, ‘dsDNA no tail’ = tailless dsDNA viruses infecting prokaryotes, ‘dsDNA tail’ = tailed dsDNA viruses infecting prokaryotes, ‘dsRNA’ = dsRNA viruses, ‘ssDNA’ = ssDNA viruses, ‘ssRNA’ = ssRNA viruses, ‘cell’ = cellular organisms (one archaeon, two bacteria, one eukaryote). Data and data sources are presented in Table S3.

We have compiled measurements of genome length and virion dimensions for 193 viruses (Table S3). Among dsDNA viruses infecting unicellular organisms, the fraction of the virion volume occupied by the viral genome declines systematically with increasing virion size (Fig. 4B), although there are notable differences among virus types. The tailless viruses infecting prokaryotes tend to have a lower fractional volume than other dsDNA viruses of the same size, and genome fractional volume declines steeply with increasing virion size for this group and for the eukaryote-infecting dsDNA viruses, which vary more than tenfold in diameter. In contrast, genome fractional volume of the tailed viruses infecting prokaryotes (members of the family *Caudovirales*) is uniformly high and weakly correlated with virion size. Regression analyses indicate that the slope of log(genome fractional volume) vs. log(equivalent spherical diameter) is ~ −1.7 for dsDNA eukaryote viruses and tailless dsDNA viruses infecting prokaryotes, while the slope for tailed phages is −0.24 (Fig. S4A). It is also noteworthy that the largest viruses overlap with small prokaryote and eukaryote cells, both in diameter and genome fractional volume, and that the largest viruses tend to infect phagotrophic eukaryotes.

A decline in genome fractional volume with virion size could be driven by selection for virion stability, because the pressure at which a capsid bursts is expected to be inversely proportional to capsid radius (Aznar et al. 2012). If dsDNA viruses generally evolve to have an internal pressure near the burst limit, then the packaging density of the genome would have to decline such that internal pressure is inversely proportional to capsid radius. However, without direct measurements it is unclear whether the observed decline in density is sufficient to equalize the stability of larger and smaller viruses, or whether larger viruses tend to be more or less stable on average. In addition, for tailed dsDNA bacteriophages the injection of the viral genome into the host cell may be driven by high genome packaging density, likely due to hydrodynamic effects of the osmotic imbalance with the host cytoplasm (Molineux and Panja 2013). In contrast, eukaryote-infecting viruses and tailless prokaryote viruses often use membrane fusion, endocytosis, or phagocytosis as an entry mechanism, although some have a more phage-like strategy (Nurmemmedov et al. 2007, Wulfmeyer et al. 2012, Mäntynen et al. 2019). Therefore, while tailed phages may require dense packaging of nucleic acids, many of the largest eukaryote-infecting viruses may reap little benefit from a pressurized capsid. The presence of a lipid envelope around the capsid may contribute to a lower genome fractional volume for many of the tailless dsDNA prokaryote viruses, but the large difference in fractional volume between these viruses and the tailed phages indicates that the envelope itself is likely not the primary cause (Fig. S4B). The dsRNA viruses may follow a scaling relationship similar to the tailless dsDNA phages, but the number of representatives in the dataset is relatively small (Fig. 4B).

Fractional genome volume also declines with virion size for ssRNA and ssDNA viruses, and these two kinds of viruses appear to follow a similar scaling relationship (Fig. 4B). This similarity is also apparent when viruses infecting multicellular organisms are also included in the comparison (Fig. S4C). The slope of the relationship is ~ −2.8 for the single-stranded viruses infecting unicells (Fig. S4A). This systematic size scaling may also be due to selection equalizing the stability of larger and smaller viruses, although the scaling relationship and mean fractional volume likely differ between single-stranded viruses and double-stranded viruses due to different physical processes underlying virion assembly and stability (Šiber et al. 2012).

Finally, physical instability may not be the primary cause of losses of infectious virions, at least in environments where solar radiation and/or non-specific adsorption are high (Suttle and Chen 1992, Noble and Furhman 1997). Viruses with larger genomes have the capacity to code for and package protective enzymes such as photolyase (Fischer et al. 2014), which could lead to slower decay rates for larger viruses. To understand the consequences of these patterns for viral fitness, and to test whether decay rate generally changes with virus size, future work should investigate decay rates in the laboratory and in natural systems for a broad size range of viruses.

### Synthesis and outlook

Here we summarize our findings on the relationships between virus size and important virus traits, and we discuss implications and future research directions.

The physics of Brownian motion predicts that smaller viruses should encounter their hosts at a faster rate (Fig. 1A), but observed adsorption rates are often much lower than theoretical contact rates (Fig. 1C), and it is possible that larger viruses have a greater attachment efficiency (Fig. 1D). New measurements of adsorption rates of large viruses are needed to test this possibility. Furthermore, entering the host cell via phagocytosis may be a particularly good strategy for ensuring that encounter leads to infection, but demonstrating this quantitatively will require studies on the mechanism of entry, and efficiency of attachment and entry, for diverse viruses.

Burst size is lower for larger viruses (Fig. 2A), which is expected if host materials and energy limit viral production, but the cost is less than would be expected if viral production was inversely proportional to genome size. This is likely due to greater control of viral replication and host physiology by viruses with larger genomes, as evidenced by their greater nucleotide production rate (Fig. 2C). Host range may generally increase with virus size, as observed for two diverse groups of phages infecting aquatic bacteria (Fig. 3), and this may be due to greater autonomy during replication, a greater range of counter-defenses, or ability to attach to a greater diversity of receptors. Finally, decay rate has not been widely studied for viruses with unicellular hosts, but a comparison of coliphages suggests that larger viruses could have a lower decay rate (Fig. 4A). In sum these observations suggest that larger viruses experience reductions in burst size, a modest or negligible reduction in adsorption rate, an increase in host range, and potentially a decline in decay rate. These countervailing selection pressures may explain how viruses of very different size have evolved and can persist when infecting the same host population.

Are there particular host traits or environmental conditions that could select for larger or smaller viruses? To address this, we can ask whether particular contexts may change the magnitude or direction of relationships between virus size and different virus traits. For example, are there conditions under which the reduction in burst size with increased virus size is lessened? The data synthesized in Figure 2 come from experiments with resource-replete host cultures, and it is possible that under host resource limitation a greater virus size is more costly (due to less energy and materials available for replication) or less costly (due to greater control of host metabolism). Testing this possibility will require measurements of the infection cycle of diverse viruses under different resource conditions. Although contact rates are predicted to decline for larger viruses due to reduced diffusivity, the magnitude of the decline is somewhat less when hosts are motile or generate feeding currents (Fig. 1A). Therefore, larger viruses may be more common among hosts that are highly motile. Likewise, if phagotrophy is an effective means of entering host cells, for viruses large enough to induce phagotrophy, then the largest viruses may be particularly prevalent among phagotrophic hosts (Fig. 4B).

If larger viruses tend to have a broader host range, the benefits of broad host range may be greatest when hosts are at low abundance. In our model of host range evolution, a virus will be able to persist if the sum of its host populations, in the absence of viral mortality, exceeds the minimum persistence threshold *S*\* (Appendix 3). Therefore, oligotrophic environments may be enriched in larger viruses if smaller viruses with narrower host ranges cannot persist. Low host density is also expected to select for lysogeny (Stewart and Levin 1984, Weitz et al. 2019), and so environments with a greater proportion of lysogenic viruses may also tend to have larger viruses in the lytic fraction. Finally, if larger size is associated with reduced decay rates then this relationship may be steeper under conditions of rapid decay, such as exposure to high insolation, which could select for larger viruses.

If we combine several of the conditions that could favor large viruses, it may be that motile, phagotrophic protists with low population densities are particularly likely to host giant viruses. Low population densities are characteristic of the oligotrophic open ocean and other environments with low nutrient or energy supply. These environments are also relatively enriched in motile, phagotrophic eukaryotes, including mixotrophic phytoplankton and heterotrophic protists (Edwards 2019), compared to more productive environments where immotile, non-phagotrophic diatoms often dominate microbial biomass. Therefore, the effects of low resource supply on both population densities and community structure may cause larger viruses to be favored in oligotrophic environments. Testing for patterns in virus size distributions across environmental conditions or host types will require a suite of methods, including substantial new isolation efforts, as well as metagenomic protocols that can capture the full size spectrum of viruses in the environment while quantitatively comparing viruses with RNA, ssDNA, and dsDNA genomes. For example, two studies compared RNA and DNA viral metagenomes in coastal ocean environments in Hawai’i (Steward et al. 2013) and Antarctica (Miranda et al. 2016) and found that the abundance of RNA viruses rivals that of DNA viruses. The RNA viruses were essentially exclusively eukaryote-infecting, while most of the DNA viruses were likely phages. This implies that RNA viruses, which tend to be smaller, are more prevalent than larger DNA viruses among eukaryotic viruses in coastal environments. No comparable studies have been performed in the open ocean, but we predict that in the pool of eukaryote-infecting viruses, larger DNA viruses, and ‘giant’ *Mimiviridae* in particular, are more prevalent than small ssDNA or ssRNA viruses in open ocean environments that tend to be more oligotrophic. It is less clear whether one should expect the size structure of prokaryote-infecting viruses to vary as much across environmental gradients in the ocean, because the abundance of prokaryotes varies much less than the abundance and biomass of unicellular eukaryotes (Li et al. 2004). A study of virus morphology across ocean regions found little variation in the structure of the bulk viral community, which is thought to be numerically dominated by bacteriophages (Brum et al. 2013). However, the locations compared in this study were open ocean environments that varied little in chlorophyll-a, a proxy for community biomass; comparing these environments to productive coastal locations may show that smaller phages become more prevalent in coastal systems.

Compilations of virus isolates across the tree of life show that bacteria and archaea are mainly infected by dsDNA viruses with a range of sizes, while eukaryote viruses primarily have RNA genomes that tend to be small, although there is a substantial minority of eukaryote-infecting DNA viruses that tend to be larger (Koonin et al. 2015, Campillo-Balderas et al. 2015). The genome composition and size distribution of eukaryote viruses may reflect alternative strategies responding to the barrier of the eukaryote nucleus, which restricts access to the host’s DNA replication and transcription machinery. Larger DNA viruses typically replicate partially or entirely in the cytoplasm, producing their own replication ‘factories’, while positive-sense single-stranded RNA viral genomes can immediately act as a template for translation (Schmid et al. 2014, Koonin et al. 2015). In our analyses of burst size and decay rate we have treated viruses with different genome types as comparable data points along the virus size spectrum (Figs. 2,4), but future work may consider whether there are important trait or niche differences between these groups that are not explained by size alone.

Finally, we have focused here on lytic viruses, because a substantial number of lytic viruses infecting unicellular aquatic organisms have been isolated and characterized. Selection for a temperate strategy of integrating into the host genome may also lead to important constraints on virus size evolution. The temperate strategy is a form of vertical symbiont transmission and is expected to be selected for when host densities are low (making horizontal transmission less likely) and when the virus presents a low fitness cost or even a benefit (Weitz et al. 2019). Considering the potential costs and benefits of prophage or provirus in the host genome, a larger virus genome will incur greater material and energy costs but will also be able to encode more functions that could benefit the host. Future work may consider how the size distribution of temperate viruses compares to lytic viruses, across environmental gradients or different host characteristics.

## Methods

### Compilation and analysis of adsorption rates

The literature was thoroughly searched for measurements of the adsorption rate of viruses that infect unicellular eukaryotes or bacteria, isolated from aquatic environments (freshwater or marine). This yielded data for seven viruses infecting heterotrophic bacteria, six viruses infecting cyanobacteria, six viruses infecting photosynthetic eukaryotes, and one virus infecting a heterotrophic flagellate (Table S1). For studies where adsorption kinetics were plotted but the adsorption rate was not reported, the rate of adsorption was estimated via regression. Three studies reported similar adsorption rates for multiple, related virus strains (Tarutani et al. 2006, Stoddard et al. 2007, Guerrero-Fereirra et al. 2011), and in these cases we averaged the rates to get a single value. One additional measurement of adsorption rate was acquired from a forthcoming study of FloV-SA1, a giant virus that infects the photosynthetic dictyochophyte *Florenciella*, isolated from the subtropical North Pacific (Schvarcz et al. *in prep*).

To compare observed adsorption rates to theoretical predictions of contact rate, the diffusion-limited contact rate was calculated from eqns. 2−3, using reported estimates of virus radius, host radius, and experimental temperature. For viruses with motile hosts we also calculated a theoretical contact rate that combines diffusion enhanced by advection (eqn. 4) and direct interception (eqn. S8). For three of the eukaryote motile hosts (*Micromonas pusilla, Heterosigma akashiwo*, and *Cafeteria roenbergensis*) estimates of swimming speed were taken from the literature (Table S1). For the eukaryote host *Florenciella*, swimming speed was arbitrarily set to 100 μm s^−1^, but other plausible values for a nanoflagellate did not significantly affect the results. For the seven motile bacterial hosts the swimming speed was set to 30 μm s^−1^, but the chosen value does not greatly affect the results because the phages had observed adsorption rates that were generally far below the theoretical contact rate even when diffusion is the only transport mechanism included (Fig. S3).

### Compilation and analysis of burst size and latent period

Compilation of burst size and latent period of phytoplankton viruses was described previously (Edwards and Steward 2018; Table S2). In brief, the literature was surveyed for studies that measured burst size and latent period of viruses isolated on phytoplankton or other microalgal hosts. Data were only included if hosts were grown in nutrient-replete media with irradiance not strongly limiting, so that experimental conditions would be as similar as possible. For this study we added data on two giant viruses, to better characterize effects of virus size for the largest viruses. These are CroV, which infects the heterotrophic flagellate *Cafeteria roenbergensis* (Tayler et al. 2018), and FloV-SA1, which infects the mixotrophic flagellate *Florenciella* (Schvarcz et al. *in prep*).

The effect of virus genome size on burst size was quantified using a generalized additive mixed model in which log10(burst size) was the response variable. Predictors included smoothers for log10(virus genome size) and log10(host genome size), and random effects for host taxon (cyanobacteria/diatom/prasinophyte/dinoflagellate/haptophyte/chlorophyte/cryptophyte/dictyochophyte/raphidophyte/pelagophyte/bicosoecid), host genus, virus type (dsDNA/dsRNA/ssDNA/ssRNA), and study (some studies reported multiple measurements on one or more strains). Similar models were constructed with virion production rate (burst size / latent period) as the response variable, and with nucleotide production rate (burst size * virus genome size / latent period) as the response variable, but these models included host growth rate as a covariate, and did not include host genome size.

### Compilation and analysis of host range data for marine phages

To test whether virus genome size is correlated with increased host range we re-analyzed data from two studies of marine phages. Sulcius and Holmfeldt (2016) reported host range (proportion of 21 host strains infected), genome size, virus family, and other information for 45 lytic phages infecting strains of *Cellulophaga baltica*, isolated from the Baltic Sea. These data were collated with information from Holmfeldt et al. (2016) classifying the phage strains into genera, in order to incorporate finer scale taxonomic categories in the statistical analysis. We then used a generalized additive mixed model to test for a relationship between phage genome size and host range. The model included a binomial error distribution, a smoother for host range, and random effects for virus family and virus genus that control for phylogenetic nonindependence in phage host range. We also included an observation-level random effect to account for potential overdispersion. A similar analysis was performed on data reported in Wichels et al. (1998), describing host range and taxonomy for 70 phages infecting bacterial strains in the genus *Pseudoaltermonas*, isolated from the North Sea. Virus family, species, and morphotype, as reported in Wichels et al. (1998), were used as random effects to control for phylogenetic non-independence.

## Acknowledgements

This work was supported by National Science Foundation awards OCE 15-59356 and RII Track-2 FED 17-36030 (to G.F.S and K.F.E.) and a Simons Foundation Investigator Award in Marine Microbial Ecology and Evolution (to K.F.E.). We thank Marcia Marston for providing information about the size of cyanophage RIM8.

## Supplemental Information

## Appendix 1. A model relating traits to fitness for a lytic virus

To consider how virus size may affect fitness we analyze a population dynamic model of a lytic virus and its microbial host, parameterized with the major quantitative traits that determine virus growth rate (Levin et al. 1977). The model is as follows:

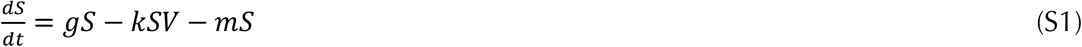

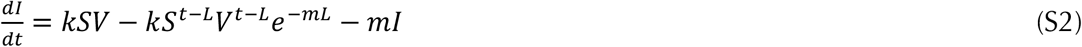

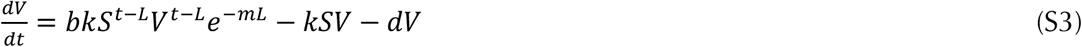

where *S* is the density of uninfected host cells, *I* is the density of infected host cells, and *V* is the density of free viral particles.

Eqn. (S1) says that the susceptible host population grows at some per capita rate *g* (the form of the growth term does not matter for this analysis), dies at a per capita rate *m*, and becomes infected at a rate *kSV*. The parameter *k* is the effective adsorption rate - the rate at which successful new infections are formed. This parameter can be decomposed into subprocesses, and here we define *k* = *caw*, where *c* is the contact rate at which host and virus encounter each other, *a* is the attachment efficiency (probability that encounter leads to successful attachment), and *w* is the probability that attachment leads to a successful infection (eventual lysis of the host, releasing new virions). The distinction between contact rate and attachment efficiency will be important when considering the role of virus size.

Eqn. (S2) says that infected cells are formed at a rate *kSV* and lost due to lysis at a rate *kS^t–L^V^t–L^e^−mL^*, where *kS^t–L^V^t–L^* is the number of cells infected at time *t - L, L* is the latent period, and *e^−mL^* is the fraction that have not perished due to other causes of mortality at rate *m*. Eqn. (S3) says that new virions are produced from lysed cells at rate *bkS^t–L^V^t–L^e^−mL^*, where *b* is the burst size (number of infectious virions produced per host cell). New virions are removed via attachment to the host at rate *kSV*, and decay at a constant rate *d*.

At steady state the density of the limiting resource, uninfected cells, is 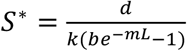 (eqn. S4). This is the density at which growth of the viral population balances the decay rate. Therefore, this is also the host density threshold required for persistence of a viral population, and if the host density is initially greater it will be cropped down to this level at steady state.

## Appendix 2. S* in a model with no latent period: competitive ability in a fluctuating environment

When modeling the impacts of lytic viruses in microbial communities the delay introduced by the viral latent period is often neglected as a simplifying tactic (Jover et al. 2013, Thingstad et al. 2014). Instead of eqns. (S1−S3) in the Appendix 1 we can write:

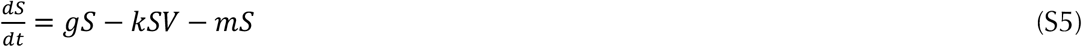

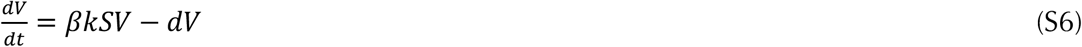

where hosts *S* are now instantly lysed, producing *β* net virions per infection. In this model the steady-state host density, at which growth of the viral population balances decay, is 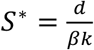 (S7). Note that the value for *S*\* derived in eqn. (S4), 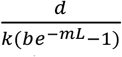, converges on 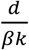 in the limit *L* → 0, setting *β* = *b* − 1 (in eqns. (S1−S3) a free virion is removed from the population when an infection is created, and this is implicitly accounted for in the value of *β*).

The dynamics described by eqns. (S5−6) may fluctuate instead of reaching a steady-state attractor, but if a second viral popuation were introduced the winner in competition would still be determined by *S*\*. This is because the per capita growth rate of a virus population, 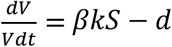, is a linear function of host density, which means that the per capita growth rate of the population over a time period *T* is:

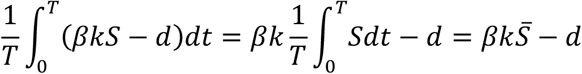

where 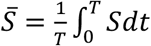 is the average host density. If the host and virus coexist then for sufficiently long *T* the virus population would neither grow forever nor decline to extinction, which means that the net population growth rate is zero. Therefore 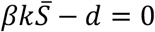 and 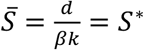. In other words the mean host density over time will be equal to the steady-state host density, and in competition the virus population with lower *S*\* will exclude its competitor by reducing the mean host density below the value its competitor requires for persistence.

## Appendix 3. Host range and competition for hosts

Virus genome size could influence host range, and so we extend the analysis of eqns. (S1-S4) to quantify how host range breadth can affect fitness. We imagine there are two host strains (A and B), each of which can be infected by a different ‘specialist’ virus strain (strains 1 and 2), and there is also a ‘generalist’ virus strain that can infect both host strains. Initially we assume the two specialists are present at steady state with no generalist – this will allow us to ask whether a broader host range could evolve. For simplicity we also assume the two specialists have the same traits, other than their host range. Under this assumption the two host strains will be cropped to the same steady state abundance:

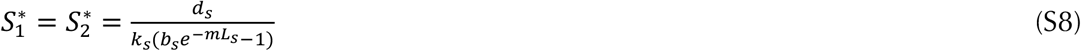

where *d*_s_, *k*_s_, *b*_s_, and *L*s are the trait values of the specialists.

Now we imagine the two host strains are not different from the perspective of the generalist (i.e., it has the same parameter values when infecting both). Then the generalist will be able to invade the two-specialist system and competitively exclude the specialists if 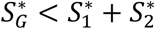, i.e., if the total abundance of the two host strains is greater than what the generalist needs to have a net zero growth rate. Alternatively if 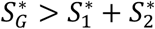 the generalist cannot invade the system. If we assume the generalist and specialists have the same decay rate then the inequality 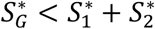 can be rearranged as follows:

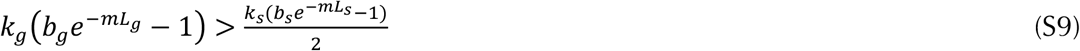

where *k*_g_, *b*_g_, and *L*_g_ are the traits of the generalist. In other words, a broader host range is favored if the burst size of the generalist is greater than half of the burst size of the specialists, or if the contact rate of the generalist is greater than half of the contact rate of the specialists, etc. This analysis could be extended to show that if a generalist has a host range that is greater than a specialist host range by a factor of *X*, it will win in competition if its fitness on each strain is at least 1/*X* times the fitness of the specialist.

This analysis of host range and competition neglects considerable complexity. For example, a generalist can coexist with a specialist under appropriate tradeoffs, and this can extend to entire networks of coexisting viruses that vary in host range (Jover et al. 2013). However, the purpose of the current analysis is simply to show that, as a first approximation, poorer competitive ability on a particular strain can be balanced by a broader host range, and the fitness benefit is (intuitively) proportional to the number of strains that can be infected.

## Appendix 4. Adsorption rate theory

An increase in diffusion-driven contact rates due to advection is quantified by the dimensionless Sherwood number (*Sh*), such that:

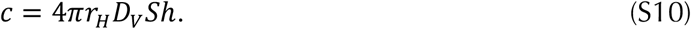

The Sherwood number can be calculated in terms of the Peclet number (Pe), a dimensionless ratio that measures the relative magnitude of advection vs. diffusion. For a sphere in motion relative to the surrounding fluid the Peclet number can be defined as 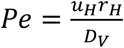, where *u_H_* is relative host velocity. An approximate formula for the Sherwood number (Murray and Jackson 1992) is then:

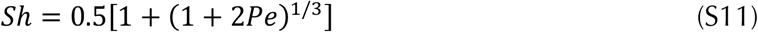

Combining eqns. S10 and S11 yields:

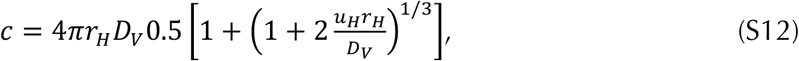

which is presented in the main text and used to plot Fig. 1A. Rates of diffusion can also be enhanced by turbulence, which will create laminar shear around a host cell at low Reynolds number. For the conditions of a virus encountering a non-motile bacterial or protist host the Sherwood number can be approximated as *Sh* = 0.955 + 0.344*Pe*^1/3^ (Karp-Boss et al. 1996).

Viruses may also encounter hosts via direct interception, where the viral particle contacts the surface of the host by occurring within the volume of water that flows within one particle radius of the cell surface, due to swimming, feeding currents, sinking, or shear (Langlois et al. 2009). For motile hosts the flow generated by swimming/feeding currents is expected to be greater than that generated by sinking or shear (Kiørboe 2008), so we focus on that source of encounter here. A simple model based on Stokes flow predicts that the contact rate is 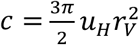 (eqn. S13; Shimeta 1993). Figure S1 compares the contact rate as a function of virus size for simple diffusion, diffusion enhanced by advection, diffusion enhanced by turbulence, and direct interception due to swimming. Diffusion driven contact tends to decline as virus size increases, although the slope of the decline is slightly less when diffusion is enhanced by advection or turbulence. Diffusion enhanced by host swimming is expected to be much more important than diffusion enhanced by turbulence (Murray and Jackson 1992). Contact via direction intercept increases as a function of virus size, but this source of encounters tends to be much less than diffusive flux, except for the largest viruses encounter hosts that swim rapidly. When swimming speed is sufficiently high the total contact rate due to diffusion and interception can exhibit a minimum near 0.5 μm virus diameter and then increase for larger viruses (Fig. S1D; Shimeta 1993). It is possible that the formula used here for direct interception underestimates the true contact rate by this mechanism (Langlois et al. 2009; others). If we arbitrarily increase the rate of interception by a factor of 3 this causes the total contact rate to bend upwards at a moderately smaller virus size (Figure S2).

**Figure S1.**
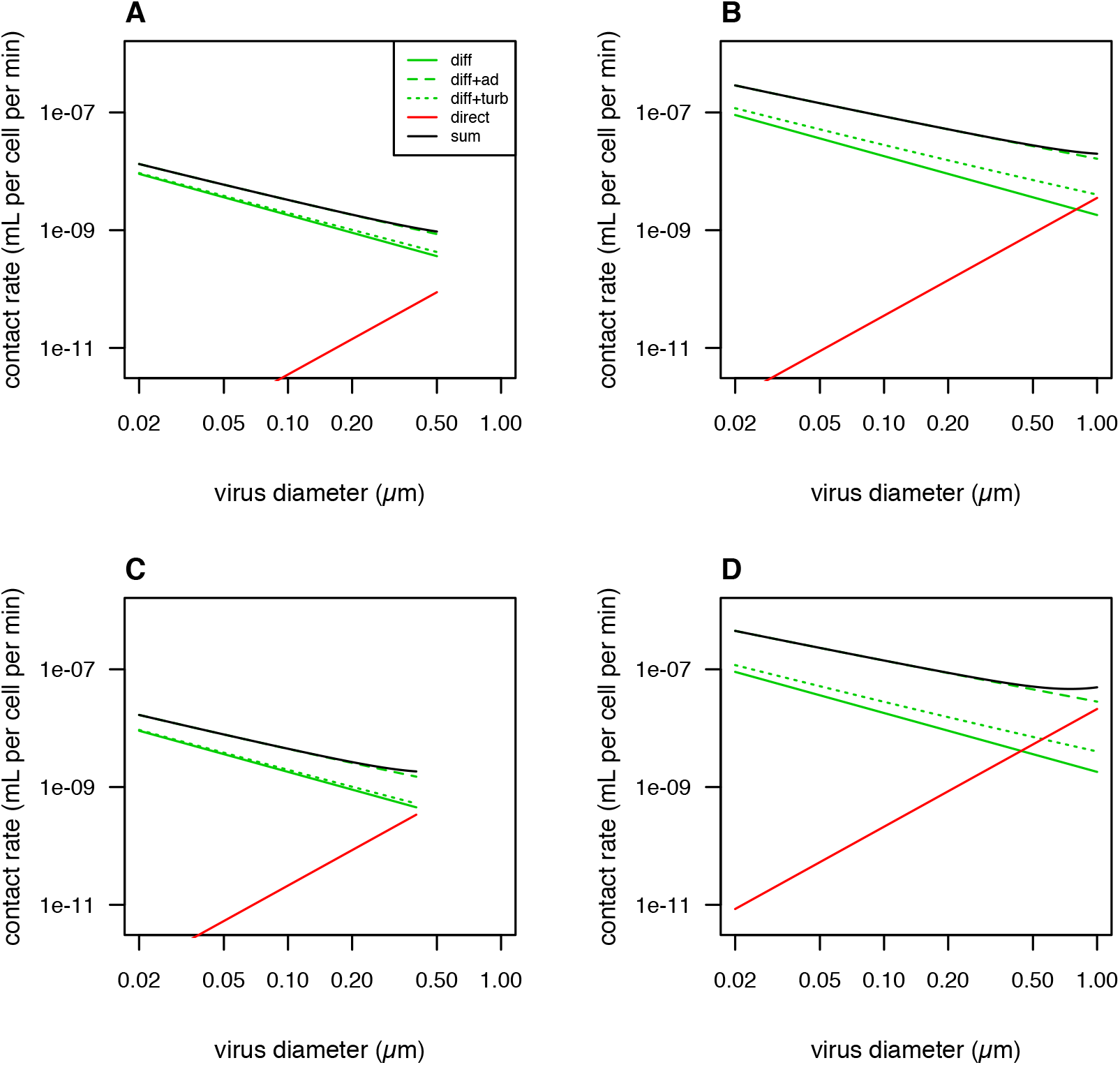
Theoretical predictions for contact rate as a function of virus size due to different physical mechanisms. ‘diff’ = only molecular diffusion, ‘diff+ad’ = diffusion enhanced by advection, ‘diff+turb’ = diffusion enhanced by turbulence, ‘direct’ = direct interception only, ‘sum’ = sum of diffusion enhanced by advection and direction interception. (A) Host diameter 1 μm, host swimming speed 5 μm s^−1^. (B) Host diameter 10 μm, host swimming speed 50 μm s^−1^. (C) Host diameter 1 μm, host swimming speed 30 μm s^−1^. (D) Host diameter 10 μm, host swimming speed 300 μm s^−1^.

**Figure S2.**
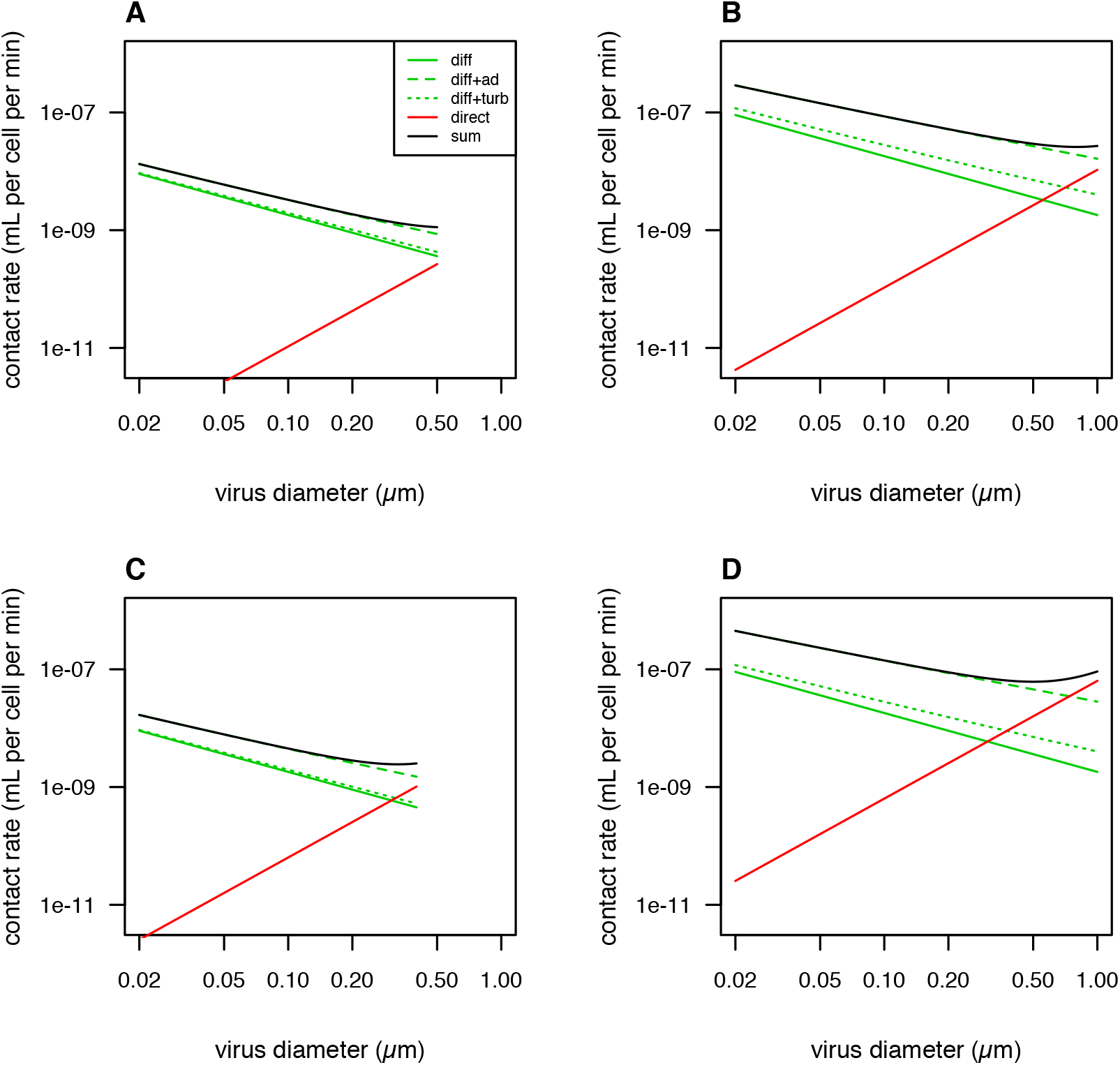
Theoretical predictions for contact rate, with rates of direct interception increased threefold. ‘diff’ = only molecular diffusion, ‘diff+ad’ = diffusion enhanced by advection, ‘diff+turb’ = diffusion enhanced by turbulence, ‘direct’ = direct interception only, ‘sum’ = sum of diffusion enhanced by advection and direction interception. (A) Host diameter 1 μm, host swimming speed 5 μm s^−1^. (B) Host diameter 10 μm, host swimming speed 50 μm s^−1^. (C) Host diameter 1 μm, host swimming speed 30 μm s^−1^. (D) Host diameter 10 μm, host swimming speed 300 μm s^−1^.

**Figure S3.**
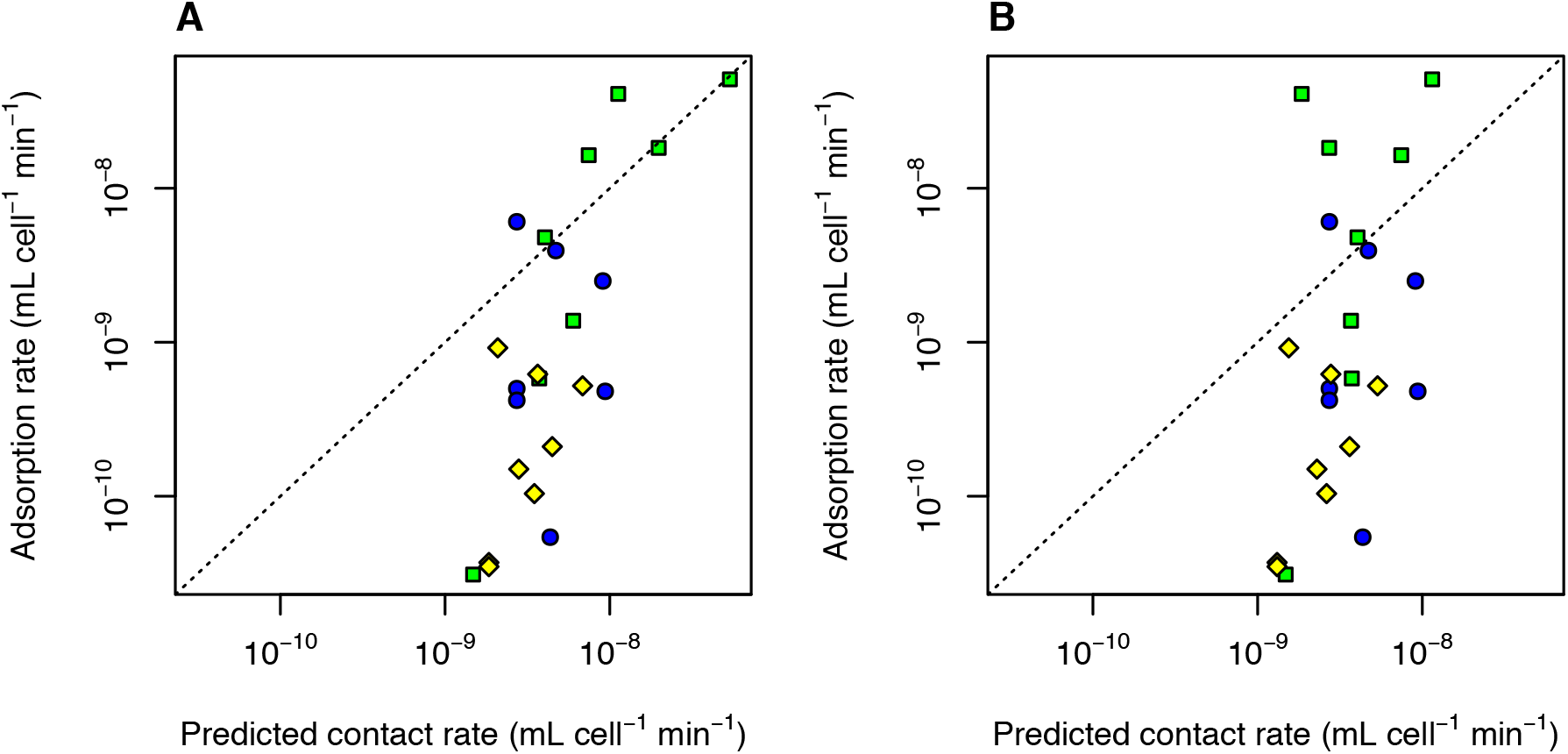
Observed adsorption rate vs. theoretical predictions of contact rate. (A) Predicted contact rate includes diffusion enhanced by advection and direct interception for motile hosts (this panel is equivalent to Fig. 1C). (B) Predicted contact rate only includes molecular diffusion, without any effects of host motility.

**Figure S4.**
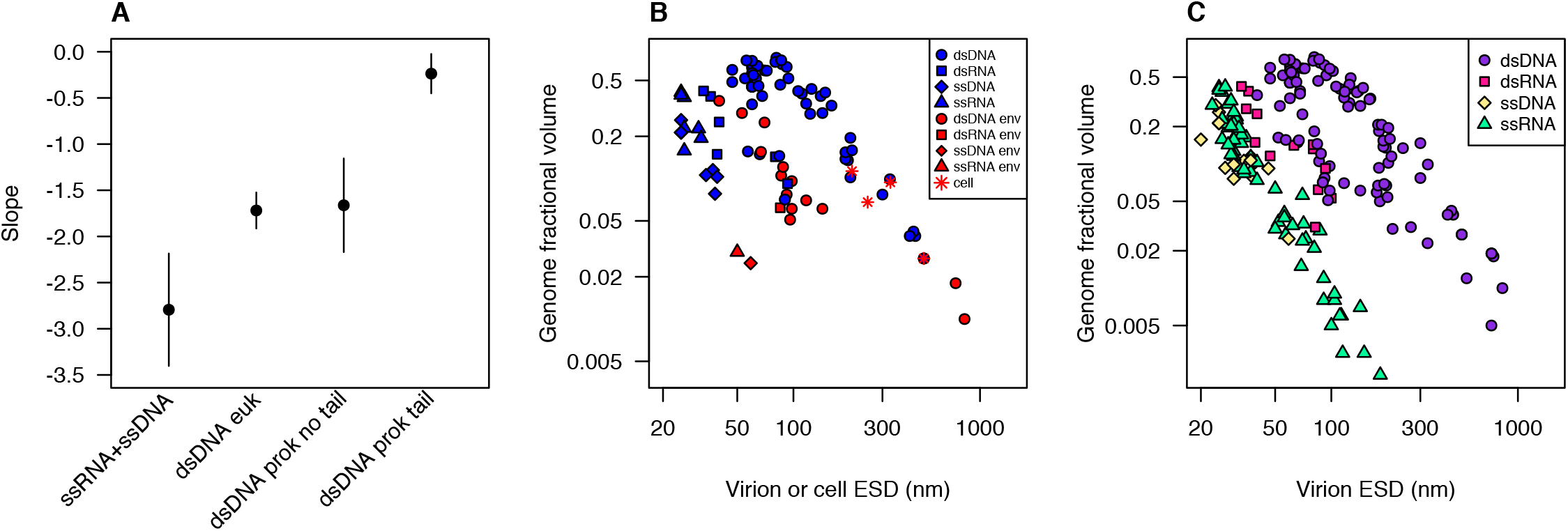
Additional analysis of genome fractional volume vs. equivalent spherical diameter. (A) Slope and 95% CI from regressions of log(genome fractional volume) vs. log(equivalent spherical diameter), for four groups of viruses infecting unicellular hosts: ssRNA and ssDNA viruses, dsDNA viruses infecting eukaryotes, tailless dsDNA viruses infecting prokaryotes, tailed dsDNA viruses infecting prokaryotes. (B) Genome fractional volume vs. equivalent spherical diameter for viruses infecting unicellular hosts, as well as four representative unicellular organisms, highlighting enveloped viruses (in red) and nonenveloped viruses (in blue). (C) Genome fractional volume vs. equivalent spherical diameter, for viruses infecting unicellular and multicellular organisms. ‘dsDNA’ = dsDNA viruses, ‘dsRNA’ = dsRNA viruses, ‘ssDNA’ = ssDNA viruses, ‘ssRNA’ = ssRNA viruses, ‘cell’ = cellular organisms (one archaeon, two bacteria, one eukaryote). Genome fractional volume is estimated nucleic acid volume divided by total virion volume. Virion volumes were estimated from reported outer dimensions (including outer envelope, if present, but excluding tails or fibrils) using volume formulae for simplified approximate shapes as indicated. Nucleic acid volume estimated as volume of a cylinder with diameter being 2.37 nm for double-stranded nucleic acid and 1.19 nm for single-stranded and 0.34 nm per nt or bp. Data and data sources are presented in Table S3.

## References

Abedon, S. T., Hyman, P., & Thomas, C. (2003). Experimental examination of bacteriophage latent-period evolution as a response to bacterial availability. Appl. Environ. Microbiol., 69(12), 7499–7506.

Al-Shayeb, B., Sachdeva, R., Chen, L.X., Ward, F., Munk, P., Devoto, A., Castelle, C.J., Olm, M.R., Bouma-Gregson, K., Amano, Y. and He, C. (2020). Clades of huge phages from across Earth’s ecosystems. Nature, 1–7.

Aznar, M., Luque, A., & Reguera, D. (2012). Relevance of capsid structure in the buckling and maturation of spherical viruses. Physical biology, 9(3), 036003.

Birch, E. W., Ruggero, N. A., & Covert, M. W. (2012). Determining host metabolic limitations on viral replication via integrated modeling and experimental perturbation. PLoS Computational Biology, 8(10), e1002746.

Brown, J. H., Gillooly, J. F., Allen, A. P., Savage, V. M., & West, G. B. (2004). Toward a metabolic theory of ecology. Ecology, 85(7), 1771–1789.

Brum, J. R., Schenck, R. O., & Sullivan, M. B. (2013). Global morphological analysis of marine viruses shows minimal regional variation and dominance of non-tailed viruses. The ISME journal, 7(9), 1738–1751.

Campillo-Balderas, J. A., Lazcano, A., & Becerra, A. (2015). Viral genome size distribution does not correlate with the antiquity of the host lineages. Frontiers in Ecology and Evolution, 3, 143.

Chow, C. E. T., & Suttle, C. A. (2015). Biogeography of viruses in the sea. Annual review of virology, 2, 41–66.

Chrzanowski, T. H., & Šimek, K. (1990). Prey-size selection by freshwater flagellated protozoa. Limnology and Oceanography, 35(7), 1429–1436.

Cui, J., Schlub, T.E. and Holmes, E.C., 2014. An allometric relationship between the genome length and virion volume of viruses. Journal of Virology, 88(11), pp.6403–6410.

De Paepe, M., & Taddei, F. (2006). Viruses’ life history: towards a mechanistic basis of a trade-off between survival and reproduction among phages. PLoS Biology, 4(7), e193.

Devoto, A.E., Santini, J.M., Olm, M.R., Anantharaman, K., Munk, P., Tung, J., Archie, E.A., Turnbaugh, P.J., Seed, K.D., Blekhman, R. and Aarestrup, F.M. (2019). Megaphages infect Prevotella and variants are widespread in gut microbiomes. Nature Microbiology, 4(4), 693.

Edwards, K. F., & Steward, G. F. (2018). Host traits drive viral life histories across phytoplankton viruses. The American Naturalist, 191(5), 566–581.

Edwards, K. F., Thomas, M. K., Klausmeier, C. A., & Litchman, E. (2012). Allometric scaling and taxonomic variation in nutrient utilization traits and maximum growth rate of phytoplankton. Limnology and Oceanography, 57(2), 554–566.

Epstein, S. S., & Shiaris, M. P. (1992). Size-selective grazing of coastal bacterioplankton by natural assemblages of pigmented flagellates, colorless flagellates, and ciliates. Microbial Ecology, 23(3), 211–225.

Felsenstein, J. (1985). Phylogenies and the comparative method. The American Naturalist, 125(1), 1–15.

Finkel, Z. V, & Irwin, A. J. (2001). Light absorption by phytoplankton and the filter amplification correction: cell size and species effects. Journal of Experimental Marine Biology and Ecology, 259(1), 51–61.

Fischer, M. G., Allen, M. J., Wilson, W. H., & Suttle, C. A. (2010). Giant virus with a remarkable complement of genes infects marine zooplankton. Proceedings of the National Academy of 5ciences, 107(45), 19508–19513.

Fischer, M. G., Kelly, I., Foster, L. J., & Suttle, C. A. (2014). The virion of Cafeteria roenbergensis virus (CroV) contains a complex suite of proteins for transcription and DNA repair. Virology, 466, 82–94.

Fuchs, H. L., & Franks, P. J. S. (2010). Plankton community properties determined by nutrients and size-selective feeding. Marine Ecology Progress Series, 413, 1–15.

Goldhill, D. H., & Turner, P. E. (2014). The evolution of life history trade-offs in viruses. Current opinion in virology, 8, 79–84.

Heineman, R. H., Springman, R., & Bull, J. J. (2008). Optimal foraging by bacteriophages through host avoidance. The American Naturalist, 171(4), E149–E157.

Heldal, M., & Bratbak, G. (1991). Production and decay of viruses in aquatic environments. Mar. Ecol. Prog. Ser, 72(3), 205–212.

Holen, D. A., & Boraas, M. E. (1991). The feeding behavior of Spumella sp. as a function of particle size: Implications for bacterial size in pelagic systems. Hydrobiologia, 220(1), 73–88.

Holmfeldt, K., Middelboe, M., Nybroe, O., & Riemann, L. (2007). Large variabilities in host strain susceptibility and phage host range govern interactions between lytic marine phages and their Flavobacterium hosts. Appl. Environ. Microbiol., 73(21), 6730–6739.

Holmfeldt, K., Solonenko, N., Howard-Varona, C., Moreno, M., Malmstrom, R. R., Blow, M. J., & Sullivan, M. B. (2016). Large-scale maps of variable infection efficiencies in aquatic Bacteroidetes phage-host model systems. Environmental Microbiology, 18(11), 3949–3961.

Holmfeldt, K., Solonenko, N., Shah, M., Corrier, K., Riemann, L., VerBerkmoes, N. C., & Sullivan, M. B. (2013). Twelve previously unknown phage genera are ubiquitous in global oceans. Proceedings of the National Academy of Sciences, 110(31), 12798–12803.

Jover, L. F., Cortez, M. H., & Weitz, J. S. (2013). Mechanisms of multi-strain coexistence in host-phage systems with nested infection networks. Journal of Theoretical Biology, 332, 65–77.

Karp-Boss, L., Boss, E., & Jumars, P. A. (1996). Nutrient fluxes to planktonic osmotrophs in the presence of fluid motion. Oceanography and Marine Biology, 34, 71–108.

Kauffman, K.M., Hussain, F.A., Yang, J., Arevalo, P., Brown, J.M., Chang, W.K., VanInsberghe, D., Elsherbini, J., Sharma, R.S., Cutler, M.B. and Kelly, L. (2018). A major lineage of non-tailed dsDNA viruses as unrecognized killers of marine bacteria. Nature, 554(7690), 118.

Kiørboe, T. (2008). A mechanistic approach to plankton ecology. Princeton University Press.

Koonin, E. V, & Yutin, N. (2019). Evolution of the large nucleocytoplasmic DNA viruses of eukaryotes and convergent origins of viral gigantism. Adv. Virus Res, 103, 167–202.

Langlois, V. J., Andersen, A., Bohr, T., Visser, A. W., & Kiørboe, T. (2009). Significance of swimming and feeding currents for nutrient uptake in osmotrophic and interception-feeding flagellates. Aquatic Microbial Ecology, 54(1), 35–44.

Le, S., He, X., Tan, Y., Huang, G., Zhang, L., Lux, R., Shi, W. and Hu, F. (2013). Mapping the tail fiber as the receptor binding protein responsible for differential host specificity of Pseudomonas aeruginosa bacteriophages PaP1 and JG004. PloS One, 8(7), e68562.

Lenski, R. E. (1988). Experimental studies of pleiotropy and epistasis in Escherichia coli. I. Variation in competitive fitness among mutants resistant to virus T4. Evolution, 42(3), 425–432.

Levin, B. R., Stewart, F. M., & Chao, L. (1977). Resource-limited growth, competition, and predation: a model and experimental studies with bacteria and bacteriophage. The American Naturalist, 111(977), 3–24.

Li, W. K., Head, E. J., & Harrison, W. G. (2004). Macroecological limits of heterotrophic bacterial abundance in the ocean. Deep Sea Research Part I: Oceanographic Research Papers, 51(11), 1529–1540.

Li, Z., Wu, J., & Wang, Z. G. (2008). Osmotic pressure and packaging structure of caged DNA. Biophysical journal, 94(3), 737–746.

Lin, T.-Y., Lo, Y.-H., Tseng, P.-W., Chang, S.-F., Lin, Y.-T., & Chen, T.-S. (2012). A T3 and T7 recombinant phage acquires efficient adsorption and a broader host range. PloS One, 7(2), e30954.

Maat, D. S., & Brussaard, C. P. D. (2016). Both phosphorus-and nitrogen limitation constrain viral proliferation in marine phytoplankton. Aquatic Microbial Ecology, 77(2), 87–97.

Mäntynen, S., Sundberg, L. R., Oksanen, H. M., & Poranen, M. M. (2019). Half a century of research on Membrane-Containing bacteriophages: bringing new concepts to modern virology. Viruses, 11(1), 76.

Miranda, J. A., Culley, A. I., Schvarcz, C. R., & Steward, G. F. (2016). RNA viruses as major contributors to Antarctic virioplankton. Environmental microbiology, 18(11), 3714–3727.

Molineux, I. J., & Panja, D. (2013). Popping the cork: mechanisms of phage genome ejection. Nature Reviews Microbiology, 11(3), 194–204.

Murray, A. G., & Jackson, G. A. (1992). Viral dynamics: a model of the effects of size, shape, motion and abundance of single-celled planktonic organisms and other particles. Marine Ecology Progress Series, 103–116.

Noble, R. T., & Fuhrman, J. A. (1997). Virus decay and its causes in coastal waters. Appl. Environ. Microbiol., 63(1), 77–83.

Nurmemmedov E., Castelnovo M., Catalano CE, Evilevitch A.. (2007). Biophysics of viral infectivity: matching genome length with capsid size. Q Rev Biophys, 40, 327–356.

Purohit, P. K., Kondev, J., & Phillips, R. (2003). Mechanics of DNA packaging in viruses. Proceedings of the National Academy of 5ciences, 100(6), 3173–3178.

Rodrigues, R. A. L., Abrahão, J. S., Drumond, B. P., & Kroon, E. G. (2016). Giants among larges: how gigantism impacts giant virus entry into amoebae. Current Opinion in Microbiology, 31, 88–93.

Samimi, B., & Drews, G. (1978). Adsorption of cyanophage AS-1 to unicellular cyanobacteria and isolation of receptor material from Anacystis nidulans. Journal of Virology, 25(1), 164–174.

Samson, J. E., Magadán, A. H., Sabri, M., & Moineau, S. (2013). Revenge of the phages: defeating bacterial defences. Nature Reviews Microbiology, 11(10), 675.

Schmid, M., Speiseder, T., Dobner, T., & Gonzalez, R. A. (2014). DNA virus replication compartments. Journal of virology, 88(3), 1404–1420.

Schwartz, M. (1976). The adsorption of coliphage lambda to its host: effect of variations in the surface density of receptor and in phage-receptor affinity. Journal of Molecular Biology, 103(3), 521–536.

Schwarzer, D., Buettner, F.F., Browning, C., Nazarov, S., Rabsch, W., Bethe, A., Oberbeck, A., Bowman, V.D., Stummeyer, K., Mühlenhoff, M. and Leiman, P.G. (2012). A multivalent adsorption apparatus explains the broad host range of phage phi92: a comprehensive genomic and structural analysis. Journal of Virology, 86(19), 10384–10398.

Sharon, I., Battchikova, N., Aro, E.M., Giglione, C., Meinnel, T., Glaser, F., Pinter, R.Y., Breitbart, M., Rohwer, F. and Béja, O. (2011). Comparative metagenomics of microbial traits within oceanic viral communities. The ISME Journal, 5(7), 1178.

Shimeta, J. (1993). Diffusional encounter of submicrometer particles and small cells by suspension feeders. Limnol. Oceanogr., 38, 456–465.

Shinn, G. L., & Bullard, B. L. (2018). Ultrastructure of Meelsvirus: A nuclear virus of arrow worms (phylum Chaetognatha) producing giant “tailed” virions. PloS One, 13(9), e0203282.

Šiber, A., Božič, A. L., & Podgornik, R. (2012). Energies and pressures in viruses: contribution of nonspecific electrostatic interactions. Physical chemistry chemical physics, 14(11), 3746–3765.

Šimek, K., & Chrzanowski, T. H. (1992). Direct and indirect evidence of size-selective grazing on pelagic bacteria by freshwater nanoflagellates. Appl. Environ. Microbiol., 58(11), 3715–3720.

Steward, G. F., Culley, A. I., Mueller, J. A., Wood-Charlson, E. M., Belcaid, M., & Poisson, G. (2013). Are we missing half of the viruses in the ocean?. The ISME journal, 7(3), 672–679.

Stoddard, L. I., Martiny, J. B. H., & Marston, M. F. (2007). Selection and characterization of cyanophage resistance in marine Synechococcus strains. Appl. Environ. Microbiol., 73(17), 5516–5522.

Storms, Z. J., & Sauvageau, D. (2015). Modeling tailed bacteriophage adsorption: Insight into mechanisms. Virology, 485, 355–362.

Šulčius, S., & Holmfeldt, K. (2016). Viruses of microorganisms in the Baltic Sea: current state of research and perspectives. Marine Biology Research, 12(2), 115–124.

Suttle, C. A., & Chen, F. (1992). Mechanisms and rates of decay of marine viruses in seawater. Appl. Environ. Microbiol., 58(11), 3721–3729.

Talmy, D., Beckett, S. J., Zhang, A. B., Taniguchi, D. A. A., Weitz, J. S., & Follows, M. J. (2019). Contrasting controls on microzooplankton grazing and viral infection of microbial prey. Front. Mar. Sci, 6, 182.

Tarutani, K., Nagasaki, K., & Yamaguchi, M. (2006). Virus adsorption process determines virus susceptibility in Heterosigma akashiwo (Raphidophyceae). Aquatic Microbial Ecology, 42(3), 209–213.

Tétart, F., Repoila, F., Monod, C., & Krisch, H. M. (1996). Bacteriophage T4 host range is expanded by duplications of a small domain of the tail fiber adhesin. Elsevier.

Thamatrakoln, K., Talmy, D., Haramaty, L., Maniscalco, C., Latham, J.R., Knowles, B., Natale, F., Coolen, M.J., Follows, M.J. and Bidle, K.D. (2019). Light regulation of coccolithophore host–virus interactions. New Phytologist, 221(3), 1289–1302.

Thingstad, T. F., Våge, S., Storesund, J. E., Sandaa, R.-A., & Giske, J. (2014). A theoretical analysis of how strain-specific viruses can control microbial species diversity. Proceedings of the National Academy of Sciences, 111(21), 7813–7818.

Thomas, R., Grimsley, N., Escande, M., Subirana, L., Derelle, E., & Moreau, H. (2011). Acquisition and maintenance of resistance to viruses in eukaryotic phytoplankton populations. Environmental Microbiology, 13(6), 1412–1420.

Tilman, D. (1982). Resource competition and community structure. Princeton university press.

Tomaru, Y., Mizumoto, H., Takao, Y., & Nagasaki, K. (2009). Co-occurrence of DNA-and RNA-viruses infecting the bloom-forming dinoflagellate, Heterocapsa circularisquama, on the Japan coast. Plankton and Benthos Research, 4(4), 129–134.

Wang, N., Dykhuizen, D. E., & Slobodkin, L. B. (1996). The evolution of phage lysis timing. Evolutionary Ecology, 10(5), 545–558.

Weitz, J. S., Li, G., Gulbudak, H., Cortez, M. H., & Whitaker, R. J. (2019). Viral invasion fitness across a continuum from lysis to latency. Virus evolution, 5(1), vez006.

Wichels, A., Biel, S. S., Gelderblom, H. R., Brinkhoff, T., Muyzer, G., & Schütt, C. (1998). Bacteriophage diversity in the North Sea. Appl. Environ. Microbiol., 64(11), 4128–4133.

Wickham, T. J., Granados, R. R., Wood, H. A., Hammer, D. A., & Shuler, M. L. (1990). General analysis of receptor-mediated viral attachment to cell surfaces. Biophysical Journal, 58(6), 1501–1516.

Wilhelm, S.W., Bird, J.T., Bonifer, K.S., Calfee, B.C., Chen, T., Coy, S.R., Gainer, P.J., Gann, E.R., Heatherly, H.T., Lee, J. and Liang, X., 2017. A student’s guide to giant viruses infecting small eukaryotes: from Acanthamoeba to Zooxanthellae. Viruses, 9(3), 46.

Wilson, W. H., Carr, N. G., & Mann, N. H. (1996). The effect of phosphate status on the kinetics of cyanophage infection in the oceanic cyanobacterium Synechococcus sp. WH7803. Journal of Phycology, 32(4), 506–516.

Wulfmeyer, T., Polzer, C., Hiepler, G., Hamacher, K., Shoeman, R., Dunigan, D.D., Van Etten, J.L., Lolicato, M., Moroni, A., Thiel, G. and Meckel, T. (2012). Structural organization of DNA in chlorella viruses. PLoS One, 7(2).

Yau, S., Caravello, G., Fonvieille, N., Desgranges, É., Moreau, H., & Grimsley, N. (2018). Rapidity of genomic adaptations to prasinovirus infection in a marine microalga. Viruses, 10(8), 441.

You, L., Suthers, P. F., & Yin, J. (2002). Effects of Escherichia coli physiology on growth of phage T7 in vivo and in silico. Journal of Bacteriology, 184(7), 1888–1894.

